# Supramolecular complexes of GCAP1: towards the development of effective biologics for inherited retinal dystrophies

**DOI:** 10.1101/2024.03.07.583919

**Authors:** A. Biasi, V. Marino, G. Dal Cortivo, D. Dell’Orco

## Abstract

Guanylate Cyclase Activating Protein 1 (GCAP1) is a neuronal Ca^2+^-sensor protein expressed in photoreceptors where it regulates the enzymatic activity of retinal Guanylate Cyclase 1 (GC1) in a Ca^2+^-dependent manner. Recently, over 20 missense mutations in *GUCA1A* (encoding for GCAP1) have been associated with inherited autosomal dominant retinal diseases, namely cone dystrophy (COD) and cone-rod dystrophy (CORD). Since GCAP1 is known to be a functional dimer, COD/CORD patients present a heterogeneous pool of GCAP1 assemblies constituted by wild-type and mutated homodimers and heterodimers. Here, we present an integrated *in silico* and biochemical investigation on the effects of the E111V substitution, associated with a severe form of CORD, on GCAP1 homo- and hetero-dimerization. Despite inducing a constitutive activation of GC1 due to impaired Ca^2+^-binding in the high-affinity EF-hand 3 motif, the E111V substitution did not affect either the homo- or the hetero dimerization process as clearly highlighted by aSEC and molecular docking experiments. Indeed, both variants exhibited micromolar monomer-dimer equilibrium constants in the presence of both Mg^2+^ and Ca^2++^, indicating that at physiological cellular concentrations both variants are predominantly monomers under Ca^2+^-loaded and, to a lesser extent, Mg^2+^-loaded conditions. Molecular docking and dynamics simulations confirmed chromatographic results highlighting slight alterations in free energy of binding involving the pathogenic E111V variant in the Ca^2+^-bound state and increased mobility over time affecting the Ca^2+^-coordinating EF3 motif. In addition, to evaluate possible therapeutic approaches, the regulation of the catalytic activity of GC1 by WT and E111V-GCAP1 was studied in the presence of retinal degeneration protein 3 (RD3), an α-helical protein that strongly inhibits GC1, and a RD3-derived peptide (RD3ppt) which encompasses a region of RD3 that is essential for its inhibitory activity. GC1 activity assays in the presence of RD3ppt suggest that the enzymatic activity is partially inhibited by the peptide at low micromolar concentrations when GCAP1 variants are present. The incomplete shut down of GC1 by RD3 could be explained by the interaction occurring between RD3 and GCAP1, known to form a complex with GC1 in the endoplasmic reticulum. This fundamental interaction was here investigated spectroscopically and *in silico*, unveiling major structural rearrangements upon complex formation. Interestingly, the full RD3 protein was able to better modulate GC1 activity and restore the abnormal cGMP production induced by the pathogenic E111V-GCAP1 variant to a physiological level.

## 1 Introduction

Visual perception occurs thanks to phototransduction, a complex biochemical cascade which ultimately converts light absorption by opsins in retinal photoreceptor cells into a change in cell membrane potential, leading to a neural response [1]. The activated opsin catalyzes the guanosine diphosphate (GDP) for guanosine triphosphate (GTP) exchange on the heterotrimeric G-protein, transducin. The GTP-bound α-subunit of transducin activates the enzyme phosphodiesterase 6 (PDE6) which in turn hydrolyzes cyclic guanosine monophosphate (cGMP) into guanosine monophosphate (GMP), decreasing intracellular cGMP concentrations. The depletion of cGMP induces the closure of cyclic nucleotide-gated (CNG) ion channels, conduits for Ca^2+^/Na+, K^+^ ions, leading to a drastic drop in Ca^2+^ concentration from several hundred nM in the dark to less than 100 nM in bright light [2]. The reduced Ca^2+^ and Na^+^ influx in photoreceptors outer segments results in cellular hyperpolarization which attenuates the release of the neurotransmitter glutamate from the synaptic terminals of the photoreceptors. The resultant modulation in glutamate release alters the activation state of the post-synaptic bipolar cells, effectively propagating the visual signal to the retinal ganglion cells and onward to the cerebral cortex. The cascade’s termination and the photoreceptor’s restoration to its basal state necessitate the deactivation of (rhod)opsins and the replenishment of cGMP. The intrinsic GTPase activity of transducin facilitates its own deactivation via GTP hydrolysis, concurrently inactivating PDE6 [1]. On the other hand, in response to diminished Ca^2+^ levels, guanylate cyclases (GCs) are activated by guanylate cyclase-activating proteins (GCAPs), thereby synthesizing cGMP anew. Maintaining calcium homeostasis is imperative for the photoreceptor’s response to light and fluctuations in intracellular calcium concentrations are sensed by GCAP proteins, which modulate the photoreceptor’s sensitivity to light. GCAPs are EF-hand neuronal Ca^2+^ sensor (NCS) proteins of the calmodulin family which are found to be active as dimers *in vitro* and to modulate cGMP synthesis in a Ca^2+^-dependent fashion by interacting with specific intracellular domains of GC [3-5]. Two GCAP isoforms, GCAP1 and GCAP2, are found in human rod and cones. The former emerges as key regulator of GC1 that represents the most impacting cyclase isozyme during the phototransduction cascade [6]. In the dark, high Ca^2+^ levels keep GCAP1 in a Ca^2+^-bound state that inhibits GC1, preventing unnecessary cGMP synthesis. Conversely, upon illumination, Ca^2+^ levels fall, prompting GCAP1 to exchange Ca^2+^ for Mg^2+^ (Figure 1A) and undergoing conformational changes that stimulate GC1 activity and rapidly replenishes cGMP, thus facilitating the restoration of the dark state and repolarization of the cell. Mutations in the *GUCA1A* gene, coding for GCAP1, have been identified as causative factors in various inherited retinal dystrophies (IRDs), such as autosomal dominant cone (COD) and cone-rod (CORD) dystrophies [7-16]. These disorders are characterized by progressive vision loss, colour vision impairment, and sensitivity to light. More than twenty pathogenic point mutations in *GUCA1A* have been identified, each leading to alterations in GCAP1’s structural and functional dynamics. The phenotypic heterogeneity observed in these dystrophies can be directly attributed to the specific amino acid changes resulting from these mutations, each affecting the protein’s ability to interact with and regulate GC1 differently, thereby disrupting the delicate equilibrium governing phototransduction. Furthermore, retinal degeneration protein 3 (RD3), a 23 kDa alpha-helical protein, has recently emerged as a key protein in the preservation and functionality of photoreceptor cells [17, 18]. RD3 steps in as a potent inhibitor maintaining a check on GCs activity with a sub-micromolar affinity and ensuring it does not prematurely engage within the photoreceptor’s inner segment, potentially averting cellular damage [19, 20]. Indeed, mutations in RD3 that affect its inhibitory activity or its binding to GC1 can lead to Leber congenital amaurosis type 12 (LCA12) and cone–rod dystrophy 6 (CORD6) [21, 22] and its absence is associated with a marked decline of GC1 and GC2 levels and their accumulation in the outer segments, implicating RD3 in the proper trafficking and stabilization of these isozymes [23, 24]. RD3s pivotal inhibitory activity arises from specific surface-exposed residues essential for the interaction with GC1 which are either located on the coil-coil domain between alpha-helix 1 and 2 (Tyr60, Trp62, and Leu63) or on alpha-helix 3 (Arg99, Arg101, and Gln102) [25]. This precise and strict regulatory behaviour could be exploited in the context of COD and CORD diseases in order to re-establish the physiological homeostasis of cGMP and Ca^2+^. The present study presents an integrated *in silico* and biochemical investigation on the effects of GCAP1 homo- and hetero-dimerization in patients carrying E111V (Figure 1B) substitution associated with a severe form of CORD. The mutation affects the high-affinity Ca^2+^ -binding motif EF3 and leads to constitutive activation of GC1 [14]. Despite inducing a constitutive activation, the E111V substitution did not affect the homo- and hetero dimerization process of GCAP1 as clearly highlighted by aSEC and molecular docking experiments, the latter highlighting subtle but severely impacting alterations of binding energies in GC1-activating and inhibiting conditions. Furthermore, to evaluate possible therapeutic approaches, the regulation of the catalytic activity of GC1 by WT and E111V-GCAP1 was studied in the presence of RD3 and RD3ppt, a RD3-derived peptide that encompasses the alpha-helix 3 region of the protein, proven to be essential for the inhibitory activity. GC1 activity assays in the presence of RD3 (Figure 6, left panel) or RD3ppt (Figure 4A) suggest that the enzymatic activity is partially inhibited when GCAP1 variants are present. The incomplete shut down of GC1 by RD3/RD3ppt reported here or by others [26] could be explained by a combination of: (i) the interaction occurring between RD3 and GCAP1, proven to be essential for the proper trafficking of the GC1-GCAP1-RD3 assembly to the photoreceptor outer segments (POS) [27]; (ii) the competitive regulation of GC1 by RD3 and GCAP1 [19]. Here the interaction between either WT- or E111V-GCAP1 and RD3/RD3ppt was investigated in GC1-activating and inhibiting conditions by CD spectroscopy experiments which underpinned major structural variations upon the RD3(ppt)-GCAP1 complex formation potentially affecting GC1 regulation and preventing rigid-body docking simulations from determining a unique binding pose. On the other hand, the full RD3 protein demonstrated increased inhibitory effect upon GC1 activity and was able to attenuate the abnormal cGMP production induced by the pathogenic E111V-GCAP1 variant, especially when an excess of WT-GCAP1 was supplied to the system, thus almost restoring the physiological regulation of GC1 and the subsequent Ca^2+^ and cGMP homeostasis.

**Figure 1.**
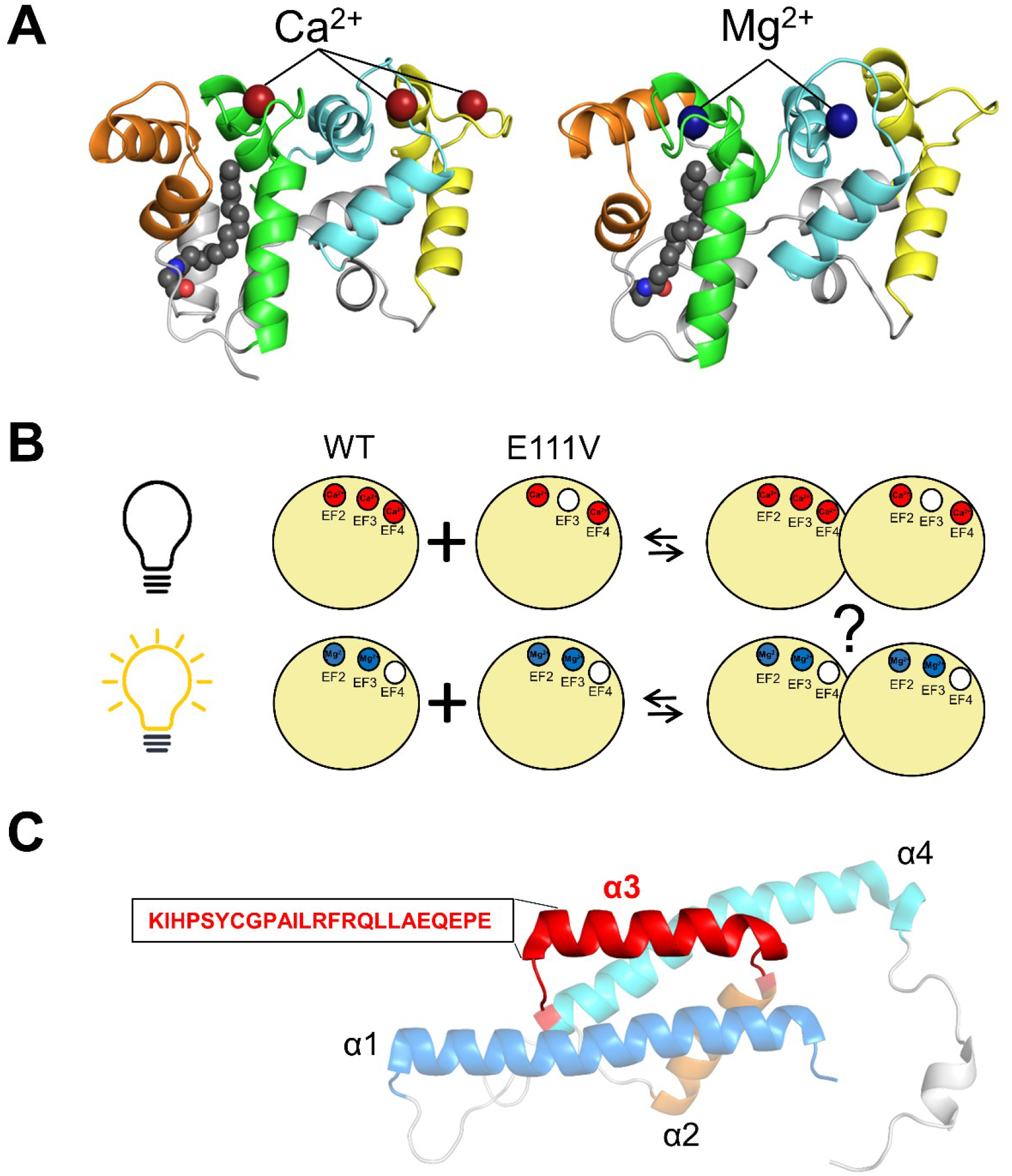
(A) Cartoon representation of the three-dimensional homology model of Ca^2+^ (left) and Mg^2+^-loaded human WT-GCAP1 in its monomeric state; EF1 is colored in orange, EF2 in green, EF3 in cyan and EF4 in yellow. N- and C-terminal are represented in light grey; Ca^2+^ and Mg^2+^ ions are shown as red and blue spheres respectively. (B) Representative diagram portraying the dimerization process investigated in this study of WT- and E111V-GCAP1 in their inhibiting (upper images) and activating states (lower images). (C) Cartoon representation of the three-dimensional NMR structure of RD3 (PDB entry 6DRF); α-helix 1 is colored in light blue, α-helix 2 in orange, α-helix 3 in red and α-helix 4 in cyan. In addition, sequence of α-helix 3 is reported and represents the RD3 peptide (RD3ppt) used.

## 2 Materials and Methods

### 2.1 Protein Expression and Purification

The pET-11a vector was utilized to clone both wild-type human GCAP1-E6S (Uniprot entry: P43080) and GCAP1-E6S-His-tag cDNAs via NdeI and NheI restriction sites. The purpose of introducing the E6S mutation was to attain a consensus sequence that could be myristoylated at the N-terminal post-translation by S. cerevisiae N-myristoyltransferase (yNMT), as supported by prior research. Subsequently, GCAP1 variants were heterologously expressed in BL21 E. Coli cells after co-transformation with pBB131-yNMT. To extract proteins from inclusion bodies, denaturation with 6 M guanidine-HCl was engaged, followed by renaturation through dialysis against 20 mM Tris-HCl pH 7.5, 150 mM NaCl, 7.2 mM β-mercaptoethanol buffer. The refolded WT- and E111V-GCAP1 variants were purified by size exclusion chromatography (SEC, HiPrep 26/60 Sephacryl S-200 HR, GE Healthcare) followed by anion exchange chromatography (AEC, HiPrep Q HP 16/10, GE Healthcare) as previously described [14]. Post-purification protein concentration was assessed by the Bradford assay [28] using a GCAP1-specific reference curve based on the amino acid hydrolysis assay (Alphalyze). Protein purity was assessed on a 15% SDS-PAGE gel. GCAP1 variants were exchanged into decalcified 50 mM NH_4_HCO_3_ buffer and lyophilized and flash-frozen in liquid nitrogen. Samples were stored at -80°C until use. Plasmid containing RD3 cDNA was a kind gift of Prof. Dott. Koch K.W. (Department of Neuroscience, Carl von Ossietzky Universität Oldenburg). RD3 was expressed in BL21 E. Coli cells and purified by a series of centrifugation steps as previously reported [19], briefly: harvested cells were mechanically lysed with 3 ultrasonication cycles (30s ON, 30s OFF). After centrifugation at 10,000 x g for 10 min, the insoluble material has been washed 3 times against 10 mM Tris-HCl pH 7.5, 2 mM EDTA, 14 mM β-mercaptoethanol, 100 µM PMSF and 1X protein inhibitor cocktail (PIC). Sample was centrifuged 15,000 x g for 15 min and the insoluble fraction was denatured overnight using the same buffer with the addition of 8 M Urea. The next day RD3 was renatured against 2 x 300 volumes of 10 mM Tris-HCl pH 7.5, 0.1 mM EDTA and 14 mM β-mercaptoethanol and centrifuged at 10,000 x g for 10 min. Supernatant containing RD3 was collected to assess protein purity via SDS PAGE and stored at -80°C with 50% v/v glycerol.

### 2.2 RD3 peptide

The RD3ppt, encompassing the region K87-E110 of RD3 that has been proven to be essential for the inhibitory activity of the protein [25], had the following sequence: KIHPSYCGPAILRFRQLLAEQEPE and was purchased by Genscript (purity >95%, checked by HPLC). The lyophilized peptide was resuspended in pure bi-distilled water at a concentration of ∼700 µM according to manufacturer instructions and stored at -80°C until use.

### 2.3 Analytical gel filtration

Analytical size exclusion chromatography (aSEC) was engaged to investigate whether the deleterious substitution E111V could alter the monomer-dimer equilibrium of GCAP1 at physiological Ca^2+^ and Mg^2+^ concentrations. Different protein dilutions ranging from 0.8 µM to 80 µM were prepared and injected (200 µL) into a Superose 12 10/300 column (GE Healthcare) previously equilibrated with a 20 mM Tris-HCl pH 7.5 buffer containing 150 mM NaCl and 0.5 mM EGTA + 1 mM Mg^2+^ or 1 mM Mg^2+^ + 0.5 mM Ca^2+^. Elution profiles were collected at 280 nm. Dissociation constants were obtained by fitting the elution volume (V_e_) to the concentration curves using equation [1] taken from [29]:

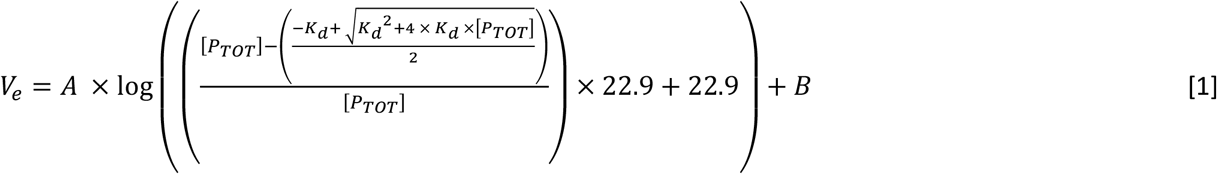

where V_e_ represents the elution volume at the peak, A is the angular coefficient, [P_TOT_] is the concentration of the protein at the time of injection, B is the y-intercept and 22.9 is the monomer theoretical M_w_ of hGCAP1. In order to accurately determine the M_w_ of eluting samples, a calibration curve has been generated by running cytochrome C (12.4 kDa), carbonic anhydrase (29 kDa), β-amylase (200 kDa), alcohol dehydrogenase (150 kDa). Consequently, the V_e_ of eluting proteins was determined by following the absorbance signal at λ = 280 nm and used to infer the distribution coefficient K_d_ using equation [2]:

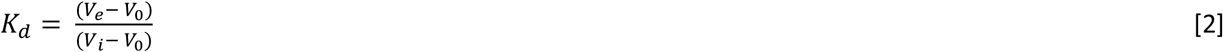

where V_0_ represents the void volume of the column (8.26 mL) and V_i_ is the total volume of the column (∼ 24 mL). Ultimately, the molecular mass (MM) of eluting proteins was determined by plotting log (MM) versus Kd.

### 2.4 Guanylate cyclase enzymatic activity assays

In this study, the effect of RD3ppt and RD3 on the regulation of GC1 activity by GCAP1 variants was investigated by performing enzymatic assays to monitor cGMP synthesis. Human recombinant ROS-GC1 was stably expressed in HEK293 cells as previously described [30]. Enzymatic experiments were performed on isolated membranes obtained after cell lysis in 10 mM HEPES pH 7.4, PIC 1X, 1 mM DTT, incubation on ice and centrifugation at 18,000 x g for 20 min and resuspended in 50 mM HEPES pH 7.4, 50 mM KCl, 20 mM NaCl and 1 mM DTT. Titrations of RD3ppt were performed with increasing concentrations of the peptide ranging from 0.05 μM to 15 μM in the presence of 5 μM WT-GCAP1 and in the almost complete absence of Ca^2+^ (∼0.073 μM). Minimum and maximum GC1 activities were determined by incubating ∼200 nM RD3 and WT or E111V-GCAP1 or both to a final concentration of 5 μM with GC1-activating buffer (2mM K_2_H_2_EGTA) or GC1-inhibiting buffer (K_2_CaEGTA) leading to <19 nM and ∼30 μM free Ca^2+^ concentrations, respectively. To assess whether RD3 affects GCAPs Ca^2+^ sensitivity (IC_50_), ∼200 nM RD3 was incubated with either WT or E111V-GCAP1 or both to a final concentration of 5 μM along with increasing free [Ca^2+^] ranging from <19 nM to 1 mM. In order to determine the GCAP1 concentration at which GC1 is active by half (EC_50_) in the presence of RD3, increasing amounts of the former (WT, E111V or both) from 0 to 15 μM were incubated with ∼200 nM of the latter in the presence of <19 nM free Ca^2+^. GC1 enzymatic reactions were performed in 30 mM MOPS/KOH pH 7.2, 60 mM KCl, 4 mM NaCl, 1 mM GTP, 3.5 mM MgCl2, 0.3 mM ATP, 0.16 mM Zaprinast buffer and blocked with the addition of 50 mM EDTA and boiling at 95°C. The synthesized cGMP was quantified by means of HPLC using a C18 reverse phase column (LiChrospher 100 RP-18, Merck). Data are reported as the mean ± standard deviation of at least three data sets.

### 2.5 Circular Dichroism

The current study employed Circular Dichroism (CD) spectroscopy on a Jasco J-710 spectropolarimeter equipped with a Peltier-type cell holder to investigate a possible interaction between GCAP1 and RD3 to unveil alterations in secondary and tertiary structure of GCAP1 upon RD3ppt or RD3 binding under different ionic conditions. GCAP1 variants were resuspended in 20 mM Tris-HCl pH 7.5, 150 mM KCl, 1 mM DTT, while the peptide and RD3 in 20 mM Tris-HCl pH 7.5, 1 mM DTT and each recorded spectrum was an average of 5 accumulations. Far-UV CD spectra of 10 µM WT-GCAP1 were acquired in a 0.1-cm quartz cuvette in the presence of 10 µM RD3ppt and after addition of 500 µM EGTA and 1 mM Mg^2+^. On the other hand, the interaction between 10 µM WT or E111V-GCAP1 and 10 µM RD3 in the far-UV was investigated after sequential addition of 300 µM EGTA and 300 µM free Ca^2+^. All CD spectra were recorded at 25 °C and normalized by mean residue ellipticity according to protein concentration and number of residues.

### 2.6 Molecular modelling, dockings and Molecular Dynamics simulations

In this work, *in silico* experiments were conducted using the Ca^2+^-loaded myristoylated GCAP1 from G. Gallus (PDB entry: 2R2I [31]) as a template, the homology model of Ca^2+^-loaded myristoylated human GCAP1 (UniProt entry: P43080) was created using the “Advanced Homology Modeling” tool offered by the software Bioluminate (Maestro package v. 12.5.139, Schrödinger). Using the “Mutate Residue” and the most likely rotamer, the E111V mutation was carried out *in silico* in the acquired human homology model. By removing the Ca^2+^ ion attached to EF-4 and replacing those in EF-2 and EF-3 with Mg^2+^, as was done in prior research [32], the activating form of human GCAP1 (Mg^2+^ -bound) was achieved. MD simulations were performed by employing the GROMACS 2020.3 package [33] and using CHARMM36m [34] as the all-atom force field which has been properly implemented with the parameters for the N-terminal myristoylated Gly (available on request). Energy minimization and equilibration procedures were engaged to prepare both GCAP1 variants in their Mg^2+^ or Ca^2+^-bound forms (2 ns in NVT ensemble with and without position restraints). Subsequently, production phase occurred and consisted of 4 independent 1 µs replicas at constant pressure (1 atm) and temperature (310 K) for each state. Resulting trajectories were used to investigate monomers flexibility by means of Root-Mean Square Fluctuation (RMSF) of the Cα which represents the time-averaged Root-Mean Square Deviation when compared to the average structure extracted by the concatenated 4 µs trajectories, following the analysis of consistency between the different replicas [35] based on the Root-Mean Square Inner Product (RMSIP) of the first 20 principal components calculated on Cα representing the largest collective motion of the protein. Docking simulations of possible dimeric assemblies were performed using ZDOCK 3.0.2 [36] and comprised 4 independent rigid-body docking runs per case tested with a sampling step of 6° (dense sampling) starting from different relative orientations each resulting in 4000 complexes. Resulting dimeric assemblies were categorized into a cohort of structurally analogous conformations, each exhibiting a Cα root-mean-square deviation (RMSD) of less than 1 Å relative to the reference complex structure [29]. Subsequently, the mean score derived from the ZDOCK algorithm, denoted as ZD, was computed for these conformationally conserved poses. Average ZD-s were utilized to approximate the binding free energy (ΔG^0^), employing the empirical relationship delineated in [37]. Moreover, consistently with CD experiments, the interaction between RD3 and GCAP1 variants was investigated *in silico*. RD3 structure was obtained by homology modelling using RD3 sequence and the 10 deposited conformers (PDB entry: 6DRF) as templates in order to obtain the full wild-type protein. In addition, the highly flexible C-terminal loop of the newly generated 10 conformers was removed. WT and E111V-GCAP1 in the activating and inhibiting conformations were docked against all 10 RD3 conformers obtained engaging ZDOCK 3.0.2 [36] and the Maestro tool PIPER [38] for a parallel analysis. Simulations with the former comprised 4 independent rigid-body docking runs, while engaging the latter up to 1000 structures have been retained probing 70000 ligand rotations in the sampling space. PIPERs retained conformations were clustered using pairwise ligand RMSD as the distance measure to identify structures at the lowest energy level. The best scoring pose among the most abundant PIPER’s output clusters was identified per each case and used as reference structure to filter ZDOCK output structures.

## 3 Results and Discussion

### 3.1 Results from MD simulations of WT/E111V-GCAP1 monomers

Previous works highlighted how the E111V substitution slightly affects secondary and tertiary structure of GCAP1 [14, 39], thus the impaired sensitivity for Ca^2+^ and the dysregulation of GC activity induced by this pathogenic variant suggested an investigation at the atomic level. 4 µs comparative MD simulations of WT and E111V-GCAP1 monomers were simulated in activating (Mg^2+^; EF2-3) and specific inhibiting conditions for each variant (Ca^2+^; WT, EF2-3-4 and E111V, EF2-(3)-4). MD trajectories revealed no significant structural perturbation during the whole simulation time, in consistency with previous spectroscopic data. However, when assessing protein structural dynamics through Cα Root-Mean Square Fluctuation (RMSF), a noticeable difference in the backbone flexibility of the E111V-GCAP1 variant was observed in comparison to the wild type (Figure 2). The mutated EF3 presents significantly higher fluctuations over time predominantly in the presence of Ca^2+^ ions. In addition, the E111V substitution primarily affects the Ca^2+^ ion allocated in EF3, promoting its detachment, and allosterically EF4 over the 4 µs of simulation time as highlighted by RMSF values calculated on the specific ion (Fig. 2B, left panel, inset).

**Figure 2.**
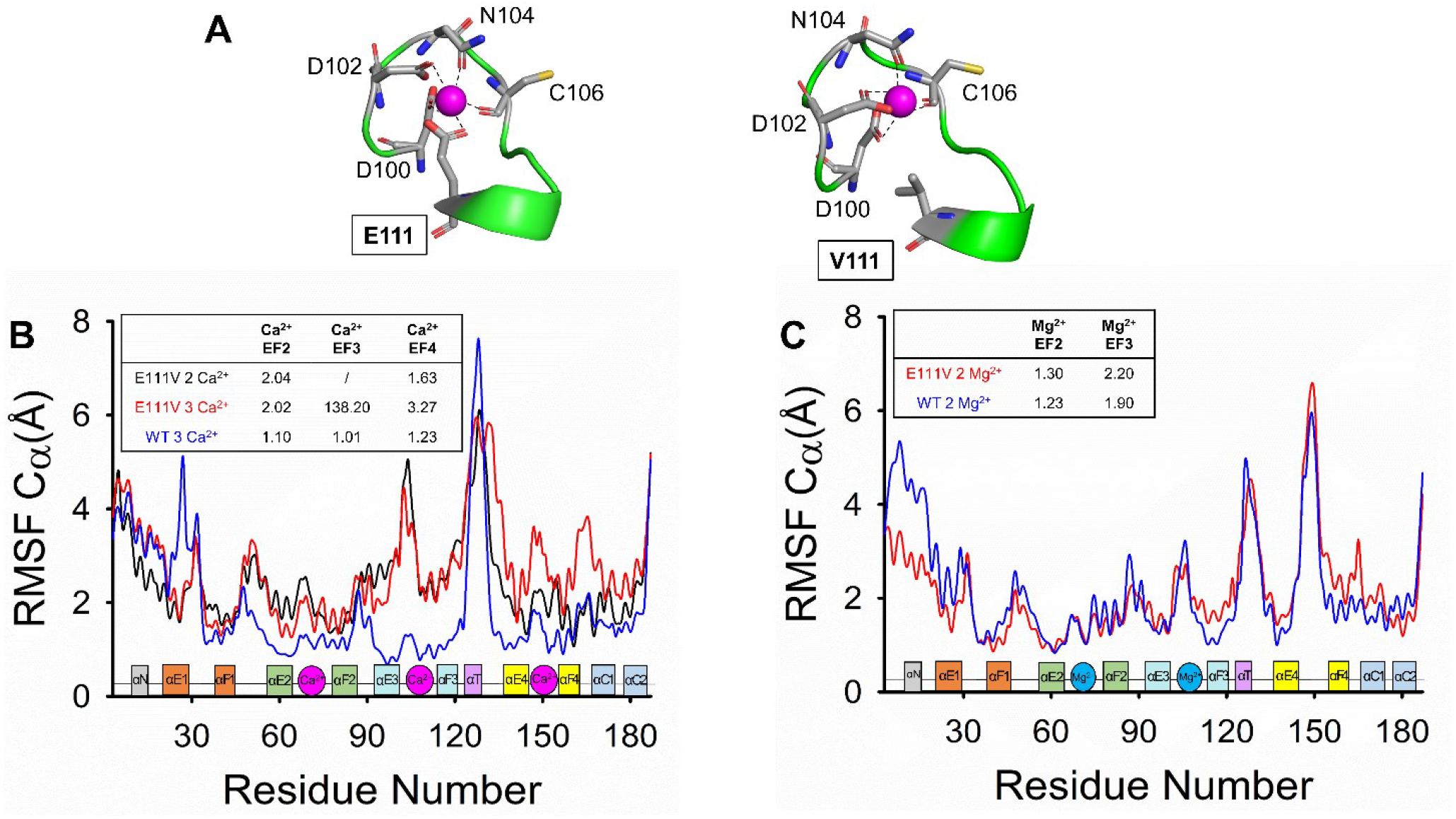
Results from exhaustive 4 µs MD simulations. (A) Representative Ca^2+^-coordination in EF3 of WT-GCAP1 (left) and E111V-GCAP1 (right) after 1 µs MD simulations. Protein structure is shown as green cartoon, Ca^2+^-coordinating residues are represented as grey sticks with O atoms in red, N atoms in blue and S atoms in yellow, Ca^2+^ ions are shown as purple spheres. (B) Cα-RMSF of Ca^2+^-loaded WT-GCAP1 (blue), E111V-GCAP1 (red), and E111V-GCAP1 with Ca^2+^-ions bound to EF2 and EF4 (red). (C) Cα-RMSF of Mg^2+^-loaded WT-GCAP1 (blue) and E111V-GCAP1 (red). Insets show the secondary structure elements colored according to Figure 1A. Inset tables show the RMSF values calculated per each ion along the 4 µs of simulation time.

### 3.2 *In silico* dimerization of WT and E111V-GCAP1 out of MD simulations

The integrity of the dimerization process, especially if driven by calcium and magnesium-bound states, is a determinant factor for GCAP1’s role in phototransduction. Hence, we investigated *in silico* whether the missense mutation could affect the dimerization process of GCAP1. Comprehensive molecular docking experiments were undertaken to elucidate the dimerization process of GCAP1 and its implications in the pathophysiology of cone dystrophies (COD) and cone-rod dystrophies (CORD), which are characterized by a pool of homo and hetero-dimers in photoreceptor cells. Rigid-body docking of potential dimeric assemblies of GCAP1 (WT/WT, E111V/E111V, WT/E111V) were tested in the same ionic conditions as MD experiments (see section 3.1) using ZDOCK software. By inferring variations in the free energy of binding (ΔΔG°), docking results highlighted subtle differences in binding affinities (Table 1); as expected the dimeric constructs exhibiting the most pronounced alterations predominantly featured the 3Ca^2+^-loaded E111V variant and to a lesser extent the 2Ca^2+^-bound form.

**Table 1.** Results from Rigid-Body Docking simulations of GCAP1 dimers. ^a^ Number of docked complexes with Cα-RMSD < 1 Å with respect to the experimentally validated dimeric model [29]; ^b^ Average RMSD of the native-like poses; ^c^ Average ZDOCK score (ZD-s) of native-like poses; ^d^ Rank of the best native-like pose out of the total 16000 proposed; ^e^ Gibbs free energy of binding; ^f^ Difference in Gibbs free energy of binding calculated with respect to WT dimers.

| Assembly | Ions | Native-like poses<sup>a</sup> | RMSD<sup>b</sup> | ZD-s<sup>c</sup> | Best ranked<sup>d</sup> | $\Delta G^{0e}$ (kcal/mol) | $\Delta\Delta G^{0f}$ (kcal/mol) |
| --- | --- | --- | --- | --- | --- | --- | --- |
| WT/WT | 2Ca | - | - | - | - | - | - |
|  | 3Ca | 22 | 0.85 ± 0.1 | 54.4 ± 0.8 | 1 | -17.36 | - |
|  | 2Mg | 22 | 0.84 ± 0.1 | 54.0 ± 0.8 | 1 | -17.20 | - |
| WT/E111V | 2Ca | 21 | 0.86 ± 0.1 | 54.6 ± 0.8 | 1 | -17.41 | -0.06 |
|  | 3Ca | 21 | 0.86 ± 0.1 | 54.8 ± 0.8 | 1 | -17.50 | -0.14 |
|  | 2Mg | 21 | 0.85 ± 0.1 | 54.3 ± 0.8 | 1 | -17.32 | -0.12 |
| E111V/E111V | 2Ca | 18 | 0.84 ± 0.1 | 54.5 ± 0.9 | 1 | -17.39 | -0.04 |
|  | 3Ca | 18 | 0.84 ± 0.1 | 54.9 ± 0.9 | 1 | -17.55 | -0.20 |
|  | 2Mg | 18 | 0.84 ± 0.1 | 54.4 ± 0.9 | 1 | -17.36 | -0.16 |

### 3.3 *In vitro* dimerization of GCAP1 variants

Molecular docking simulations highlighted neglectable differences in binding affinities upon homo and hetero-dimers formation in the presence of either Ca^2+^ or Mg^2+^ ions. Notably, despite the pathological overstimulation of GC1 by the E111V mutation, which is known to induce photoreceptor cell death, the equilibrium between monomer and dimer forms of both wild-type GCAP1 and its E111V variant at physiological conditions remains unaltered also *in vitro*. This is evidenced by analytical size exclusion chromatography (Figure 3), where both variants display similar elution profiles and dissociation constants at the equilibrium (Table 2), suggesting that the monomer-dimer transition, a critical aspect of GCAP1’s function, is resilient to the structural perturbations caused by the E111V mutation. Interestingly, at physiological cellular concentrations (3.3 μM in bovine rods, [40]), the two variants were found to be predominantly monomers under Ca^2+^-saturating conditions and, to a lesser extent, in the Mg^2+^ -bound form (Table 1). These observations underscore the specificity of the E111V mutation’s effect on GC1 stimulation, dissociating it from the fundamental dimerization capability of GCAP1, which appears to be retained.

**Table 2.** Results from CD spectroscopy. Secondary structure content estimated from CD spectra by deconvolution with BeStSel software [41]; ^a^ calculated as (θ_222_^ion^–θ_222_^EGTA^)/θ_222_^EGTA^.

| Variant | α-helix (%) | β-sheet (%) | Other (%) | $\theta_{222}/\theta_{208}$ | $\Delta\theta/\theta$ (%)<sup>a</sup> |
| --- | --- | --- | --- | --- | --- |
| RD3 | 27.7 | 12.7 | 59.6 | - | - |
| **Apo** |  |  |  |  |  |
| WT-GCAP1 | 28.3 | 16.6 | 55.1 | 0.89 | - |
| E111V-GCAP1 | 33.4 | 9.6 | 57 | 0.89 | - |
| RD3 + WT-GCAP1 | 12.4 | 29 | 58.6 | 0.98 | - |
| RD3 + E111V-GCAP1 | 13.5 | 29.4 | 57.1 | 0.99 | - |
| **Ca<sup>2+</sup>-bound** |  |  |  |  |  |
| WT-GCAP1 | 29.3 | 15.2 | 55.5 | 0.92 | 3.02 |
| E111V-GCAP1 | 32.7 | 12.6 | 54.7 | 0.90 | 2.6 |
| RD3 + WT-GCAP1 | 12.4 | 30.8 | 56.8 | 0.99 | 1.23 |
| RD3 + E111V-GCAP1 | 12.9 | 29.4 | 57.7 | 1.07 | 3.44 |

**Figure 3.**
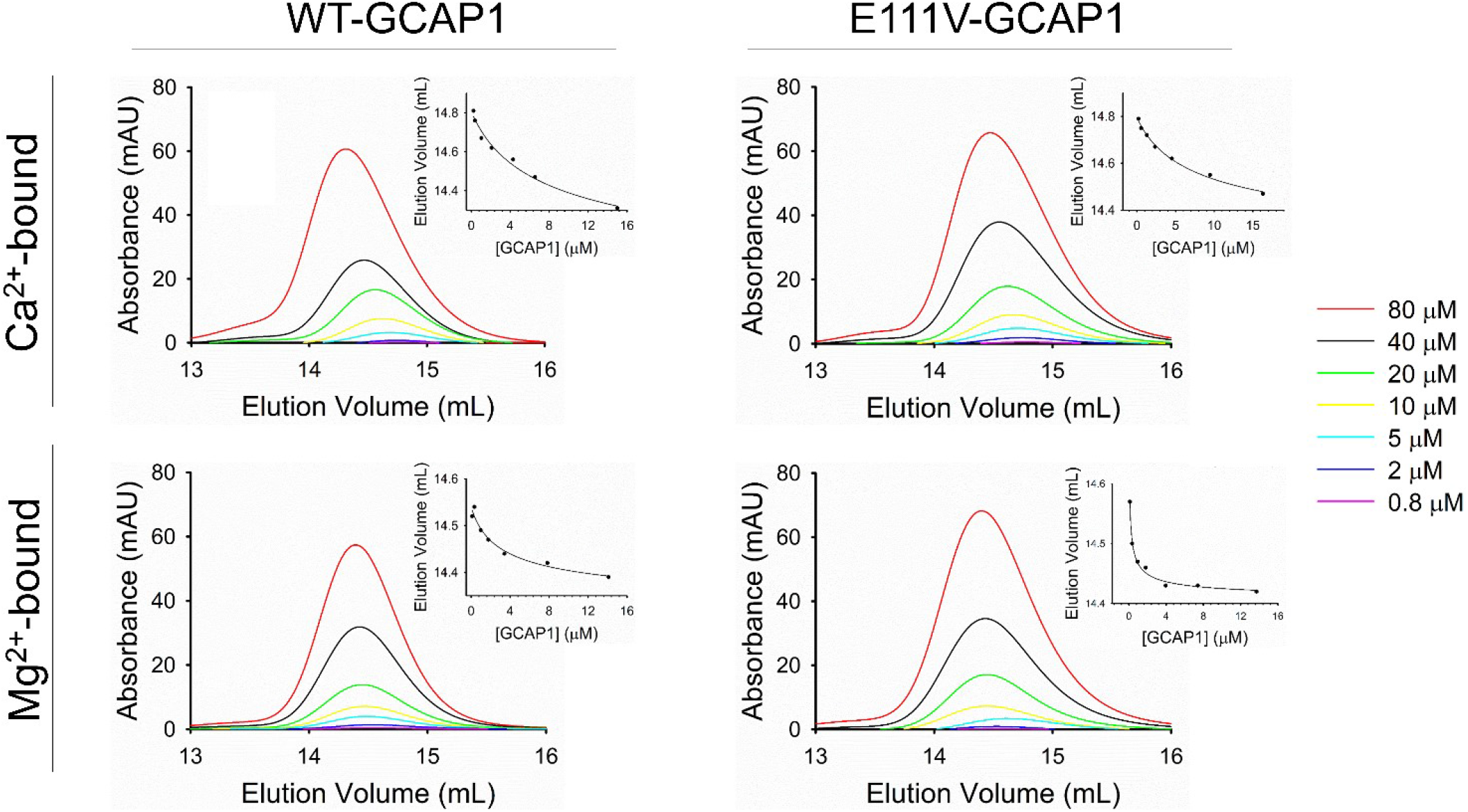
Analytical SEC chromatograms of 80 µM (red), 40 µM (black), 20 µM (green), 10 µM (yellow), 5 µM (cyan), 2 µM (blue), and 0.8 µM (purple) WT-GCAP1 (left panels) or E111V-GCAP1 (right panels) in the presence of 1 mM Mg^2+^ and 0.5 mM Ca^2+^ (upper panels) or 0.5 mM EGTA and 1 mM Mg^2+^ (lower panels). Insets show the elution volume as a function of the protein concentration at the peak together with the theoretical fitting curve (Equation 1) detailed in section 2.3.

### 3.4 RD3 inhibition of GC1: α helix 3 and whole protein

The therapeutic modulation of retinal guanylate cyclase activity constitutes a promising avenue for addressing dysregulated cGMP production implicated in IRDs. This study also focuses on the regulation of GC1 by wild-type GCAP1 and its pathogenic E111V variant in the presence of RD3 and a derived peptide, RD3ppt. The latter was observed to exert partial inhibitory control over GC1 activity at low micromolar concentrations in the presence of either WT-GCAP1 (Figure 4, panel A) or E111V-GCAP1 (Figure S1). On the other hand, the full-length RD3 protein demonstrated enhanced regulatory efficacy when titrated in the presence of GC1 and WT-GCAP1 (Figure 4, panel B). However, the incomplete shut-down of the kinase activity detected here (Figure 6, left panel) and elsewhere [24][19] could be attributed to the direct interaction occurring between GCAP1 and RD3, an interplay still poorly understood that appears to play a major role in regulating GC1 activity and viability [27]. Hence, we investigated spectroscopically and by rigid-body docking the interaction occurring between recombinant human RD3 or RD3ppt and GCAP1 variants to elucidate the molecular fingerprints defining the RD3-GCAP1 complex. CD experiments of either WT or E111V-GCAP1 in the apo and Ca^2+^-bound form highlight an interaction between GCAP1 and RD3/RD3ppt leading to a noticeable attenuation in the signal spectrum compared to individual protein measurements (Figure 5). Indeed, analysis of CD spectra underline a marked reduction in α-helical content in GCAP1 upon interacting with RD3 (Table 2), suggesting a new regulatory mechanism for the control over GC1 activity. Despite employing two distinct docking algorithms, achieving a definitive binding pose for the GCAP1-RD3 interaction proved challenging, likely due to substantial conformational changes upon complex formation, as hinted by the CD findings. In striking contrast with the GCAP1 dimerization, a flexible docking approach might offer deeper insights into this complex interaction, although it poses its own set of challenges. Remarkably, RD3 substantially moderated the aberrant cGMP synthesis instigated by the E111V-GCAP1 mutation especially when complemented with an excess of the WT form (Figure 6, Table 3). Even though the aberrant E111V activity can be effectively modulated by RD3, GC1’s activity partial inhibition suggested that its interaction with GCAP1 could also play a major role in regulating the kinase activity.

**Table 3.** Results from enzymatic assays. ^a^ Human GC1 activity as a function of free [Ca^2+^] in the presence of 5 μM GCAP1 variants and 200 nM RD3; ^b^ Hill coefficient; ^c^ GCAP1 concentration at which GC1 activity is half-maximal; ^d^ fold change in cGMP production calculated as (GC_max_ - GC_min_)/GC_min_; ^e^ Data from [39]. Data are reported as mean ± standard deviation of three technical replicates.

| | Variant | IC<sub>50</sub><sup>a</sup> ($\mu M$) | h<sup>b</sup> | EC<sub>50</sub><sup>c</sup> ($\mu M$) | X-fold<sup>d</sup> |
| --- | --- | --- | --- | --- | --- |
| - RD3 | WT<sup>e</sup> | 0.32 ± 21.6 | 2.14 ± 0.27 | 1.88 ± 0.22 | 26 |
| | E111V<sup>e</sup> | 20.2 ± 20.53 | 0.60 ± 0.38 | 1.55 ± 0.21 | 1.1 |
| | 3xWT/E111V | 8.49 ± 6.05 | 0.68 ± 0.28 | - | 3.3 |
| + RD3 | WT | 0.22 ± 0.01 | 2.23 ± 0.28 | 6.36 ± 1.17 | 14.4 |
| | E111V | 118 ± 44.8 | 0.67 ± 0.95 | 5.62 ± 1.61 | 0.76 |
| | 3xWT/E111V | 0.29 ± 0.05 | 2.24 ± 0.94 | - | 4.6 |

**Figure 4.**
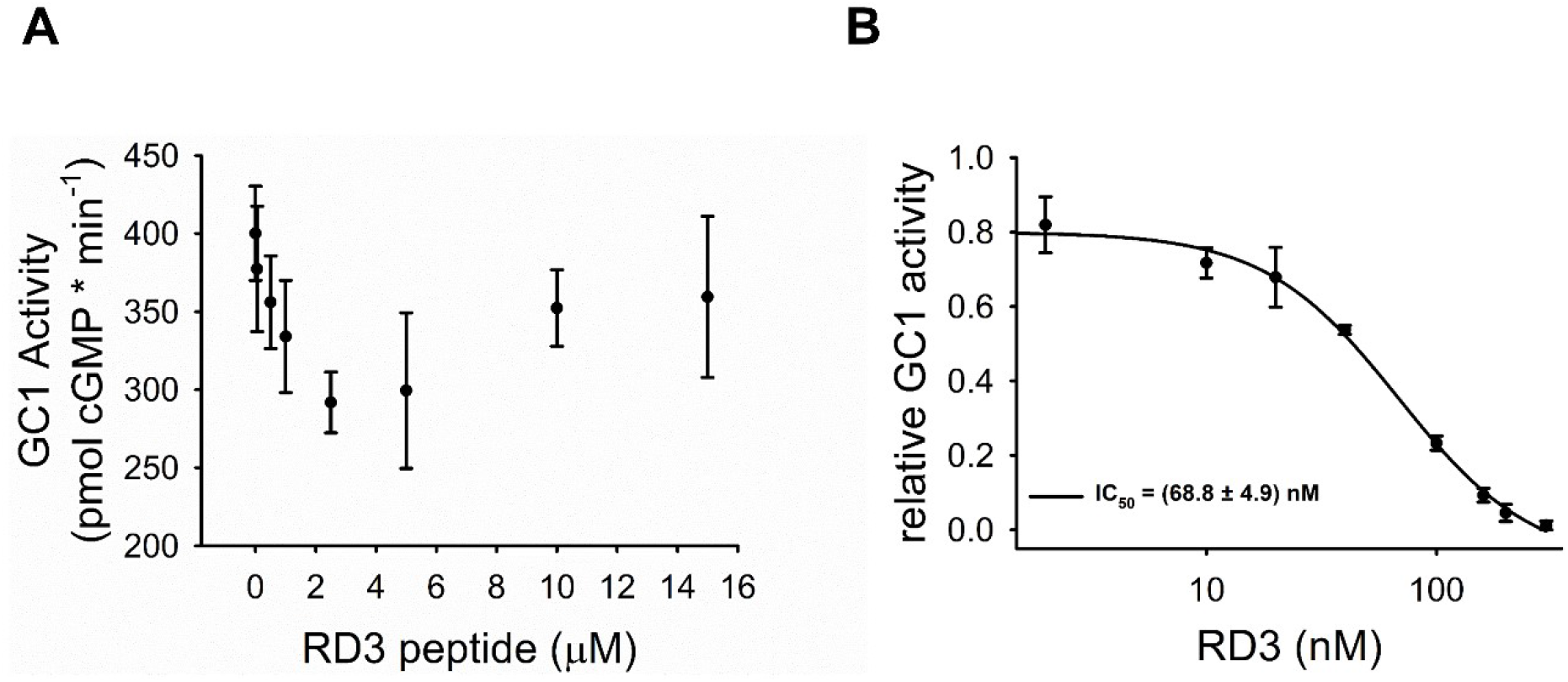
Inhibition of RetGC1: (A) the activity of recombinant RetGC1 reconstituted with 5μM purified WT-GCAP1 (black dots, mean ± S.D., n=3) was assayed at different micromolar concentrations of the RD3ppt and normalized over time. (B) Membranes containing GC1 were reconstituted with 5μM WT-GCAP1, less than 1μM free Ca^2+^ and increasing concentrations of RD3 (0-300nM). Each point is representative for 3 replicas and data are reported as relative GC1 activity and S.D. Solid black line represents the results fitting using a 4-parameter Hill equation.

**Figure 5.**
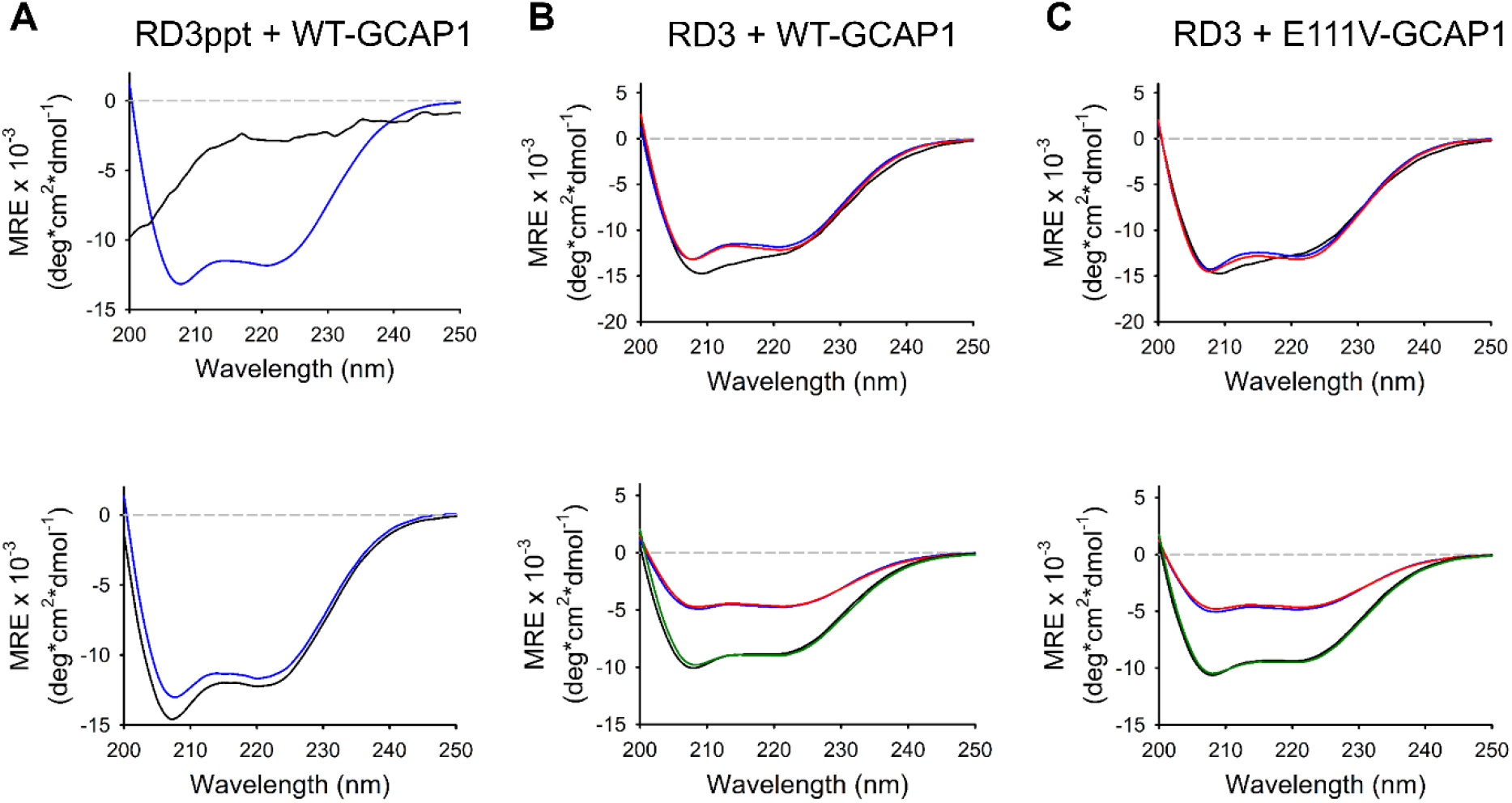
Structural changes occurring in GCAP1 variants upon RD3ppt (A) or RD3 (B and C) addition. Far UV CD spectra of: (A) (upper panel) 10 μM WT-GCAP1 (blue curve) or 10 μM RD3ppt (black curve) and (lower panel) 10 μM WT-GCAP1 + RD3ppt (blue curve) or the arithmetic sum of the single spectra of the two species (black curve) in the presence of 500 µM EGTA and 1 mM Mg^2+^; (B) (upper panel) 10 μM WT-GCAP1 or RD3 and (lower panel) 10 μM WT-GCAP1 + RD3 or the arithmetic sum of the single spectra of the two species in the presence of 300 µM EGTA (blue and black curves) and 600 µM free Ca^2+^ (red curves); (C) same as B but in the presence of 10 μM of the E111V variant. Data are normalized according to mean residue ellipticity (MRE).

**Figure 6.**
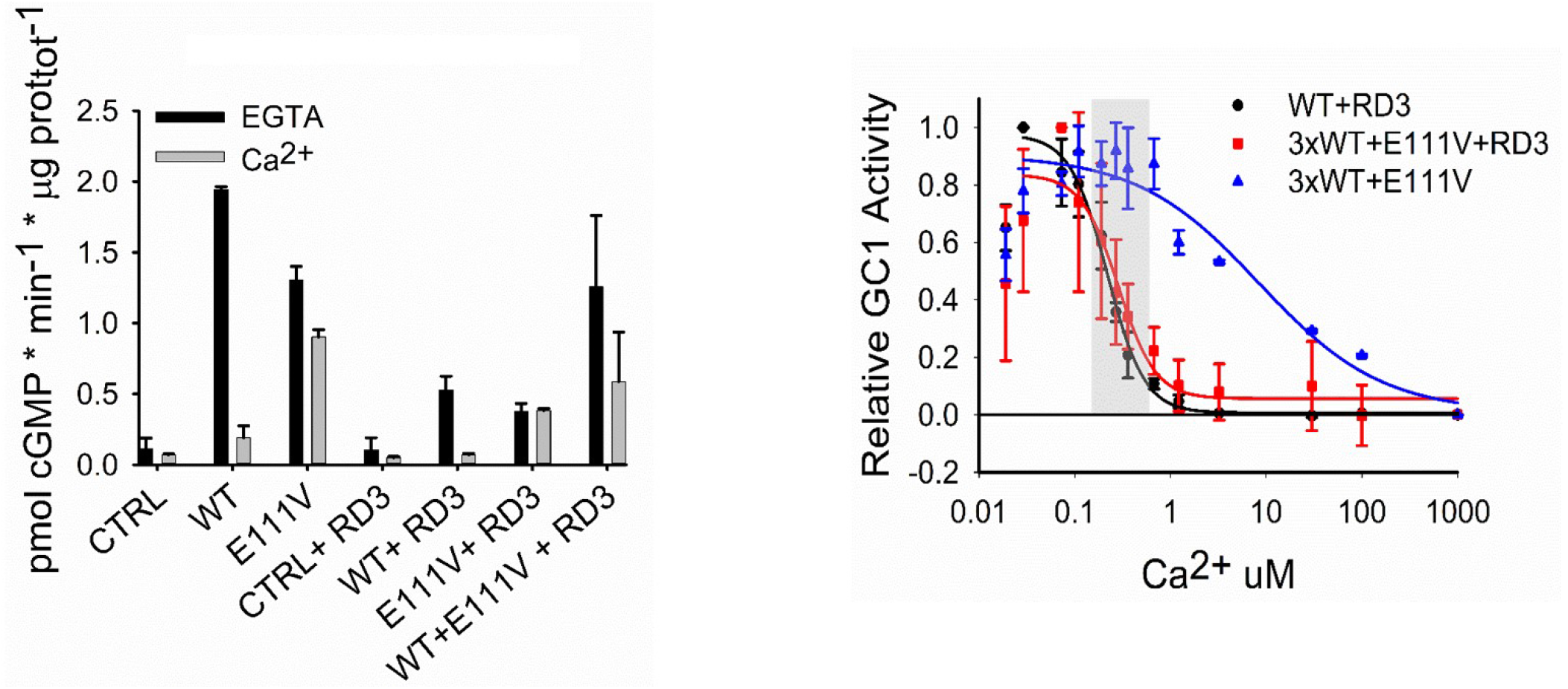
GC1 regulation by WT and E111V-GCAP1 in the absence and in the presence of 200 nM RD3. (**A**) Membranes containing GC were reconstituted with 5μM WT or E111V-GCAP1 or both and <19nM Ca^2+^ (black) or ∼30μM free Ca^2+^ (grey); membranes with no GCAP1 were used as control. (**B**) GC1 activity as function of [Ca^2+^] was measured in the presence of 5 μM WT-GCAP1 (black circles), 3.75 μM WT-GCAP1 + 1.25 μM E111V-GCAP1 + 200 nM RD3 (red squares) and 3.75 μM WT-GCAP1 + 1.25 μM E111V-GCAP1 (blue triangles). Each data set is relative to 3 replicas and data are normalized according to both total protein content in membranes and maximum and minimum GC1 activity recorded in each replica; error bars represent S.D. Solid lines represent the results of data fitting while the grey box represents the physiological Ca^2+^ fluctuations in rod photoreceptors.

## Conclusions

Based on the comprehensive research presented, several key conclusions can be drawn regarding the behaviour and interaction of GCAP1 variants and their influence on phototransduction, particularly in the context of cone-rod dystrophies (CORD). Firstly, despite the E111V mutation in GCAP1 leading to an abnormal stimulation of GC1 and subsequent photoreceptor cell death, it does not disrupt the fundamental monomer-dimer equilibrium of GCAP1. Both wild-type and E111V variants of GCAP1 displayed similar dimerization behaviour in analytical size exclusion chromatography, indicating that the dimerization process is robust against the structural changes induced by the E111V mutation. This finding is critical as it suggests that the dimerization capability of GCAP1, essential for its function in phototransduction, remains intact even in the presence of one deleterious mutation. Moreover, molecular docking experiments provided additional insights into the dimerization process of GCAP1 and its implications in photoreceptor diseases. The results showed subtle variations in the binding affinities of different dimeric constructs of GCAP1, with the most noticeable differences involving the E111V variant. This suggests that while the thermodynamics of dimerization are preserved, the E111V mutation induces specific alterations at the atomic level, affecting GCAP1’s interaction and regulation properties. The study also explored the regulatory influence of RD3 in co-presence with GCAP1 on GC1 activity. Firstly, the interaction between GCAP1 and RD3 detected *in vivo* and, here, *in vitro*, introduces a new layer of complexity in the regulation of GC1, potentially revealing previously unexplored regulatory mechanisms involving RD3 within the phototransduction pathway. Enzymatic assays highlighted RD3’s capability to mitigate the aberrant cGMP synthesis caused by the E111V mutation, especially when combined with an excess of wild-type GCAP1. This suggests a potential for RD3 to re-establish near-physiological levels of GC1 activity *in vitro*, thereby contributing to the restoration of Ca^2+^ and cGMP equilibrium disrupted in photoreceptor pathophysiology. Put together, these findings highlight the intricate interplay of GC1 regulation and present RD3, especially in its intact form, as a potent modulator capable of counterbalancing the pathological hyperactivation induced by the E111V-GCAP1 variant.

## Acknowledgments

The Centro Piattaforme Tecnologiche of the University of Verona is acknowledged for providing access to the computational and spectroscopic platforms.

## Notes

### Competing Interest Statement

The authors have declared no competing interest.

### Summary of Updates

Images and tables layouts have been revised.

