## Supplementary material for "Supramolecular complexes of GCAP1: towards the development of effective biologics for inherited retinal dystrophies": Figure S1

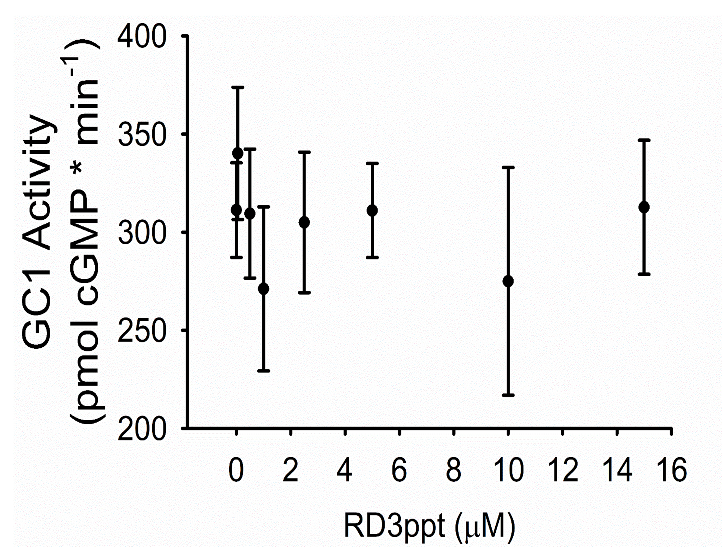

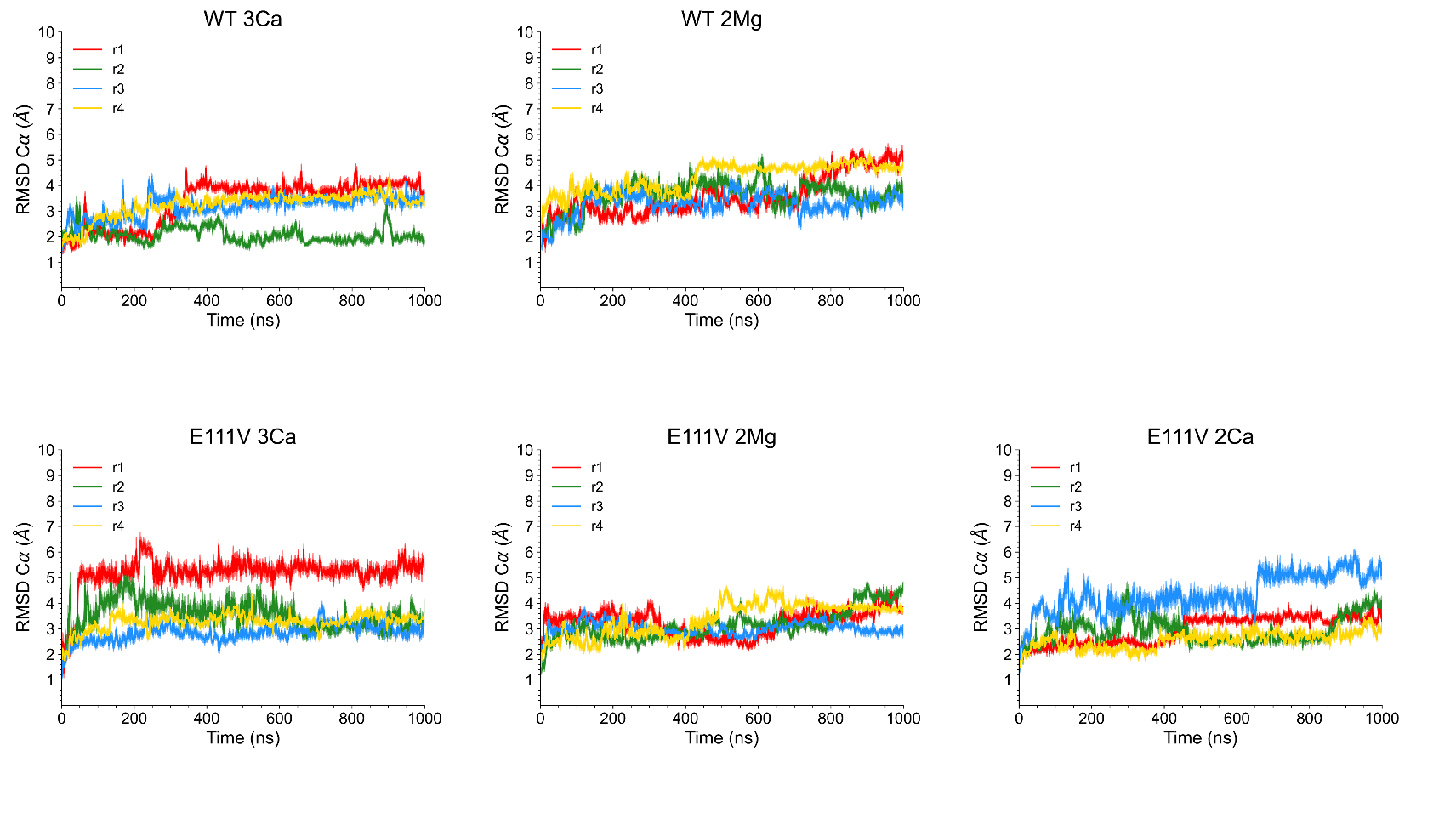
**Figure S1**. Inhibition of RetGC1 by RD3 peptide; the activity of recombinant RetGC1 reconstituted with 5μM purified E111V-GCAP1 (black dots, mean ± S.D., n=3, right panel) was assayed at different micromolar concentrations of the RD3 peptide and normalized over time.

**Figure S2**. RMSD profiles calculated per each 1 µs replica of MD simulations: (Upper panels) WT-GCAP1 and (Lower panels) E111V-GCAP1 in Mg^2+^-bound and Ca^2+^-loaded states.

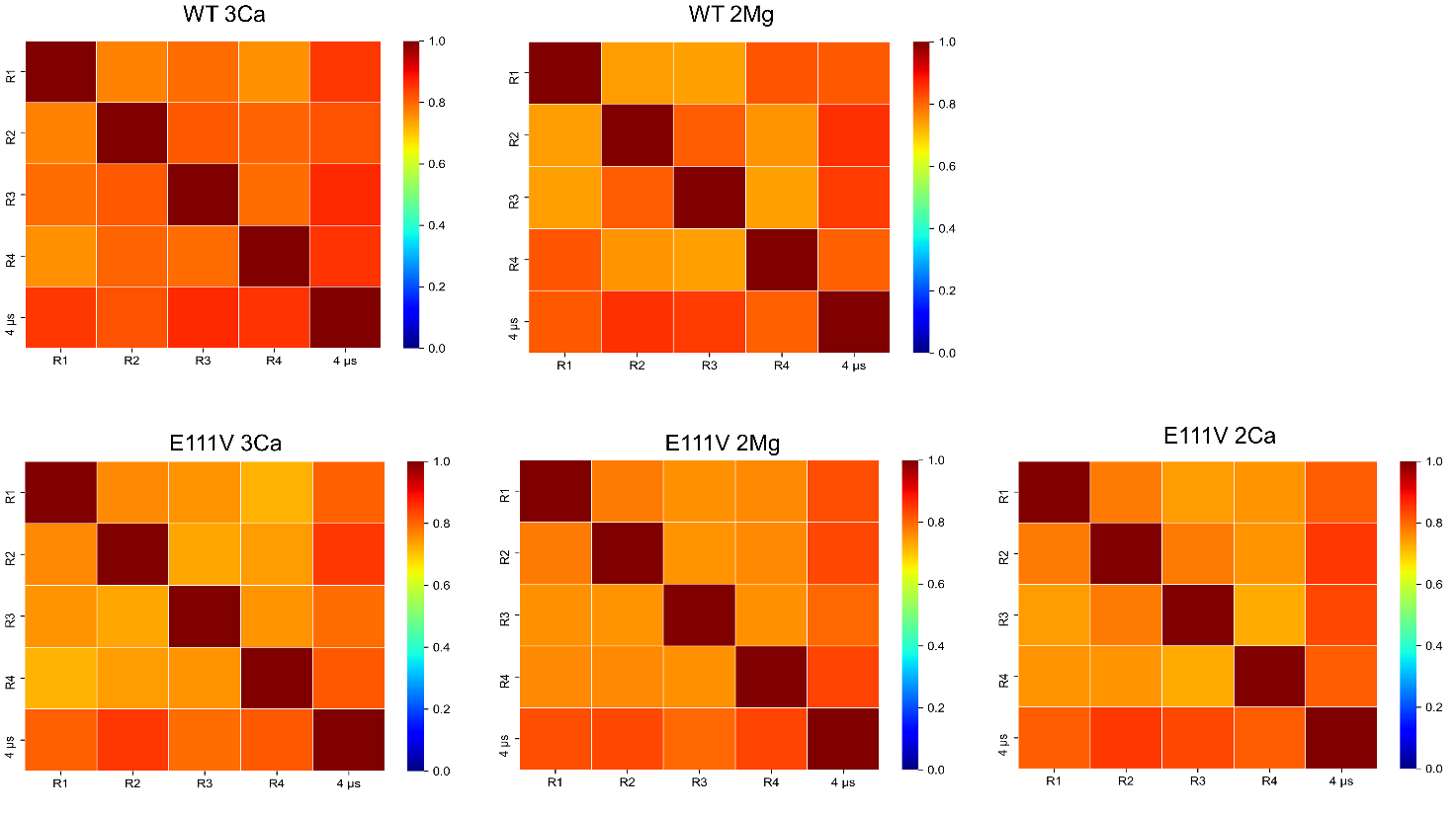

**Figure S3**. RMSIP of the first 20 principal components of the four 1 µs MD simulation replicas (R1- R4) and of the concatenated trajectories of WT- and E111V-GCAP1 in their Mg^2+^-bound and Ca^2+^-loaded forms.
